# CHARACTERIZATION OF RECOMBINASE ACTIVITY ACROSS CELLULAR GROWTH PHASES

**DOI:** 10.1101/2024.11.08.622590

**Authors:** M. Gonzalez-Colell, J. Macia

## Abstract

Recombinases, which are enzymes that catalyze targeted DNA modifications, hold significant potential in synthetic biology. Their capacity to precisely manipulate genetic material enables the construction of complex genetic circuits that can be dynamically reconfigured in response to environmental stimuli. Such capabilities are essential for developing synthetic organisms tailored for specific functions, including biosensing, bioremediation, and pharmaceutical production. Therefore, characterizing the dynamics of recombinases is crucial for the innovative design of cellular devices. A deeper understanding of how recombinases interact with DNA in various conditions can improve the efficiency and control of genetic modifications, thereby enhancing both the functionality and reliability of synthetic biological systems.

This study presents a detailed examination of the dynamics and efficiency of the serine recombinase Bxb1, focusing on its behavior under controlled expression in *Escherichia coli*. It highlights the significant influence of cellular growth phases (exponential and stationary) on the efficiency of recombinase-mediated gene expression. Our findings show that recombinase activity is maintained during stationary phase, which is critical to ensure ongoing recombination without the need for the continuous presence of an inducer.

In experiments, we quantified the recombination efficiency of Bxb1 by monitoring expression of green fluorescent protein (GFP) as a reporter. Optimal expression of Bxb1, which maximized the recombination efficiency, occurred during exponential phase. However, once the culture reached stationary phase, accumulated Bxb1 continued to facilitate recombination, although GFP expression levels plateaued due to reduced cellular activity.

These insights are relevant for synthetic biology applications where precise control of genetic functions is necessary.

## INTRODUCTION

Recombinases are enzymes that mediate the specific reorganization of DNA by binding to unique recombination sites. These sequences serve as binding points where recombinases modify DNA segments, ensuring that recombination is precise and controlled^1,2^.

In both biomedicine and synthetic biology, recombinases are valuable tools due to their ability to induce targeted and controlled genetic modifications, making them ideal for a range of applications from gene therapy to the creation of advanced biological systems. For instance, in biomedicine, recombinases are pivotal for developing innovative gene therapy strategies by enabling the insertion, deletion, or correction of genes within specific genomes, and thus have significant potential for treating genetic disorders through the direct amendment of pathogenic mutations in patients’ DNA^3,4^. Moreover, these enzymes aid the creation of transgenic animal models^5,6,7^, which provide essential tools to study human diseases and assess new therapies in vivo^8,9,10^.

In synthetic biology, recombinases are very useful for designing complex genetic circuits that regulate gene expression in response to environmental signals^11,12,13^. Furthermore, the development of recombinase-based biosensors opens up opportunities for environmental monitoring and medical diagnostics, allowing the precise detection of contaminants or disease biomarkers^14,15,16^. Moreover, the application of these enzymes in biocomputing systems marks a significant advance toward cellular devices that can be automatically activated and regulated in response to physiological signals^17,18,19,20^.

Therefore, understanding the dynamics of recombinases is critical to optimally develop cellular devices and other biotechnological or biomedical applications where recombination efficiency is key.

Focusing on synthetic biology, many applications leverage serine recombinases^21,22^, which are noted for their high specificity toward recombination sites and superior efficiency compared with other recombinases. Their flexibility means they can be used in various cell types and they exhibit low toxicity and can operate without co-factors, simplifying the design and construction of genetic circuits^22^.

This study examined the dynamics of the serine recombinase Bxb1 from bacteriophage Bxb1^23,24^, which was chosen because of its widespread use in synthetic biology. This recombinase was expressed in *Escherichia coli* under the control of the arabinose-dependent P_BAD_ promoter^25^, which allowed Bxb1 levels to be regulated based on the concentration of arabinose added to the culture.

Depending on the relative orientation of the recognition sites, i.e., *attP* and *attB*, Bxb1 can mediate two distinct types of modifications: it excises the DNA fragment flanked by the *attP* and *attB* sequences when both sequences are in the same orientation and flips this fragment when the sequences are in opposite orientations^24^. This study assessed the recombination efficiency of excision events, i.e., *attP* and *attB* sequences in the same orientation, considering two critical parameters, namely, the expression level of Bxb1 and the growth phase of the bacterial culture, focusing on exponential and stationary phases.

## RESULTS

To study the dynamics of recombination processes under different scenarios, three types of cells were constructed (Figure 1, see Supplementary Material for details). The first cell type (C1 in Figure 1) contained two plasmids. The first plasmid was the low-copy number plasmid pSB3K3^26^ which contained the gene encoding Bxb1 downstream of the arabinose-dependent promoter P_BAD_. This promoter is characterized by very low basal expression, which, combined with the low copy number of pSB3K3, ensured that basal expression of Bxb1 was minimal. This effectively prevented the occurrence of unintended recombination in the absence of arabinose.

**Figure 1.**
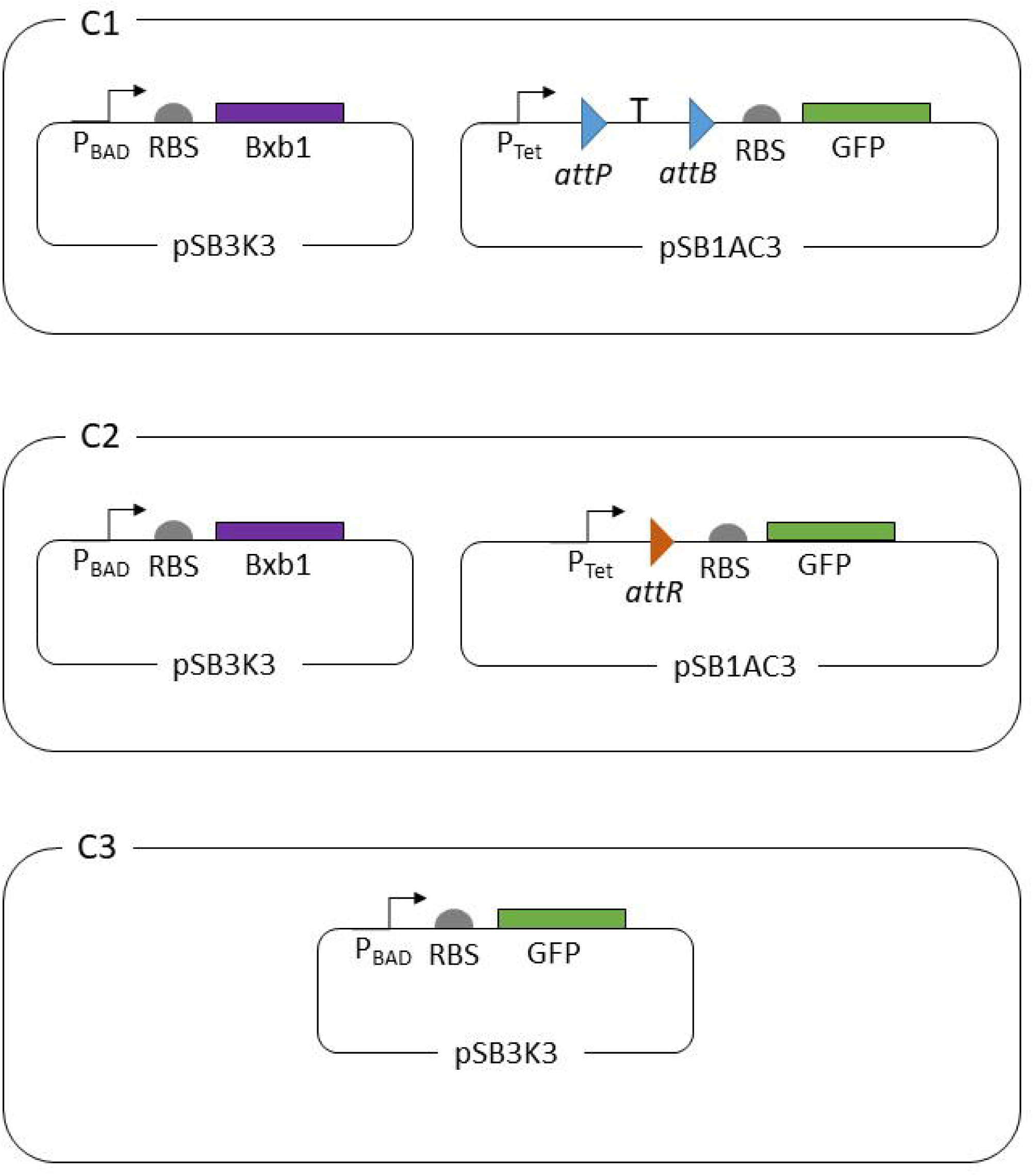
Schematic representation of the cell types used in this study. Cell type C1 contains two plasmids. The first is the low-copy number plasmid pSB3K3, which contains the gene encoding Bxb1 downstream of the arabinose-dependent promoter P_BAD_. The second is the high-copy number plasmid pSB1AC3, which contains the gene encoding GFP downstream of the strong promoter P_Tet_. However, the terminator sequence T flanked by the recombination sites *attR* and *attP* is located between the promoter and coding region, preventing gene expression. Cell type C2 is similar to cell type C1, but the terminator sequence flanked by the recombination sites is replaced by the *attR* sequence, which arises upon excision of the terminator sequence by Bxb1. Finally, cell type C3 contains the plasmid pSB3K3, in which GFP expression is controlled by the arabinose-dependent promoter P_BAD_.

The second plasmid was the high-copy number plasmid pSB1AC3^26^, which contained the gene encoding green fluorescent protein (GFP). This gene was regulated by the P_Tet_ promoter, which acts as a constitutive promoter in the absence of the TetR repressor^27^. However, a double terminator sequence was introduced between the promoter and ribosome-binding site (RBS) sequence. This terminator sequence was flanked by the recombination sites *attP* and *attB* oriented in the same direction. The presence of this terminator sequence prevented expression of GFP. In the presence of arabinose, Bxb1 was expressed, which mediated excision of the terminator sequence in this genetic construct. Consequently, GFP was expressed, allowing this plasmid to be used as a monitoring system for recombination events.

The second cell type (C2 in Figure 1) contained two plasmids. It contained the same pSB3K3 plasmid as cell type C1 together with the pSB1AC3 plasmid, which contained the GFP gene under the control of the same P_Tet_ promoter, but with the *attR* sequence introduced between the promoter and RBS. This sequence is formed as a result of the excision process^28^. Although expression of GFP was independent of Bxb1 in this cell type, it was used as a reference for the maximum fluorescence level achievable if the recombination efficiency was 100%, meaning all possible recombination sites had been modified by Bxb1.

Finally, the third cell type (C3 in Figure 1) contained the low-copy number plasmid pSB3K3 with the GFP gene downstream of the arabinose-dependent promoter P_BAD_. This cell type allowed the monitoring of P_BAD_ promoter activity in response to arabinose depending on the cell culture phase.

Using cell types C1 and C2, it was possible to quantify the recombination efficiency, denoted as ρ, mediated by Bxb1 by measuring the fluorescence levels of GFP according to the following equations:

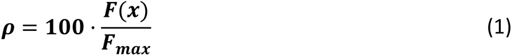

with

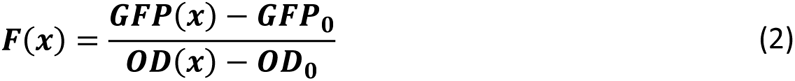

where ***GPF***(***x***) is the fluorescence produced in cell type C1 with a certain concentration of arabinose x, ***GPF***_**0**_ is the background fluorescence, ***OD***(***x***) is the optical density (OD) measured at wavelength of 600 nm of the cell culture, and ***OD***_**0**_ is the background OD measured at the same wavelength, and

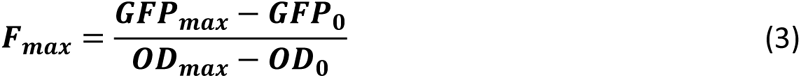

where ***GPF***_***max***_ is the fluorescence produced in cell type C2 and ***OD***_***max***_ is the OD at 600 nm of this cell culture.

### Effects of the culture growth phase on plasmid gene expression

Previous studies indicate that expression of genes carried in plasmids is highly dependent on the growth phase of the cell culture^29,30^. Specifically, gene expression is optimal during exponential phase and significantly decreases when the culture enters stationary phase. Consequently, the growth phase of the cell culture can significantly impact the outcome of recombination events due to changes in expression of Bxb1 and also of genes regulated by recombination events. To assess the impact of the growth phase of the cell culture on gene expression, cell type C3 was used, in which GFP was expressed in the same way as Bxb1 in cell type C1. A cell culture of C3 using lysogenic broth (LB) medium without arabinose was prepared and incubated at 37°C with shaking overnight. From this culture, a series of new cultures were prepared by mixing the overnight culture with fresh LB medium in various proportions (see Material and Methods for details). Specifically, the initial optical densities (OD) of the prepared cultures were 1.4, 1.25, 1.0, 0.75, 0.5, 0.25, 0.15, and 0.01, respectively. To each of these cultures, 10^-4^ M arabinose was added to induce expression of GFP. Experimental results indicate that GFP expression levels were strongly dependent on the initial concentration of the culture (Figure S1). It is worth mentioning that the duration of exponential phase varied depending on the initial concentration of the culture; higher initial culture concentrations corresponded to shorter exponential phases. Figure S2 presents the duration of exponential phase as a function of the initial culture concentration. Once all cultures reached stationary phase, GFP expression levels depended on the duration of exponential phase; shorter exponential phases corresponded to lower GFP expression levels.

According to these results, in response to an external input, e.g., arabinose, Bxb1 is produced only during exponential phase. This implies that depending on the initial culture concentration, the pool of accumulated intracellular Bxb1 varies upon exposure to the same concentration of the external input, which potentially affects the recombination efficiency. Furthermore, gene expression resulting from excision events also occurs only during exponential phase. This suggests that a reporter protein might not accurately reflect the recombination efficiency. Excision events may occur that do not have sufficient time to induce expression of the reporter protein, e.g., GFP, before stationary phase entry. Consequently, the parameter ρ in equation (1) might be underestimated. The recombination efficiency and reporter expression may be less correlated the shorter the duration of exponential phase.

Subsequently, the effects of the level of Bxb1 expression and the duration of exponential phase on resultant gene expression due to recombination events were characterized.

### Effect of the Bxb1 expression level on the recombination efficiency

To characterize the effect of Bxb1 expression levels on the efficiency of recombination events, cell type C1 (Figure 1) was used, in which Bxb1 expression was induced for different durations. Specifically, from a saturated culture of cell type C1 without arabinose, a 1:1000 dilution was prepared using LB medium containing 10^-4^ M arabinose (see Material and Methods for details). Every 30 min, a 1 mL sample of the culture was taken and centrifuged to separate the pellet from the supernatant. The pellet was resuspended in the same volume of fresh medium lacking arabinose. Finally, 1 µL of the resuspended pellet was used to inoculate 5 mL of LB lacking arabinose, and the cultures were grown overnight at 37°C with shaking. Using this procedure, a set of cultures in which the initial inoculums had been exposed to arabinose for various durations was prepared. The level of intracellular Bxb1 differed in each culture, and the recombination efficiency dependent on the duration of Bxb1 induction was determined. The final dilution of 1 µL of inoculum in 5 mL of medium lacking arabinose ensured that the duration of exponential phase was sufficiently long not to be a limiting factor.

To estimate the efficiency parameter ρ, a culture of cell type C2 was also prepared following the same procedure. Although GFP expression in cell type C2 was not dependent on Bxb1, the same procedure described for cell type C1 was performed to investigate the effects that induced Bxb1 expression might have, e.g., in terms of the metabolic burden, on GFP expression.

Figure 2a shows the relationship between Bxb1 induction time and GFP expression (orange dots). The blue dots represent GFP expression in cell type C2 and correspond to the maximum level of GFP expression achieved with 100% recombination of all possible sites. In the relationship between Bxb1 induction time and GFP expression, we distinguished three distinct behaviors. At 0–2.5 h (Region I), no significant GFP expression associated with Bxb1-mediated recombination was observed. Between 2.5 and <10 h (Region II), there was a linear relationship (r^2^=0.97) between Bxb1 induction time and GFP expression (black line in Figure 2a). Finally, at >10 h onward (Region III), 100% of recombination events had occurred and consequently maintaining Bxb1 induction beyond this time point did not elicit any additional effect on GFP expression (Figure 2b).

**Figure 2.**
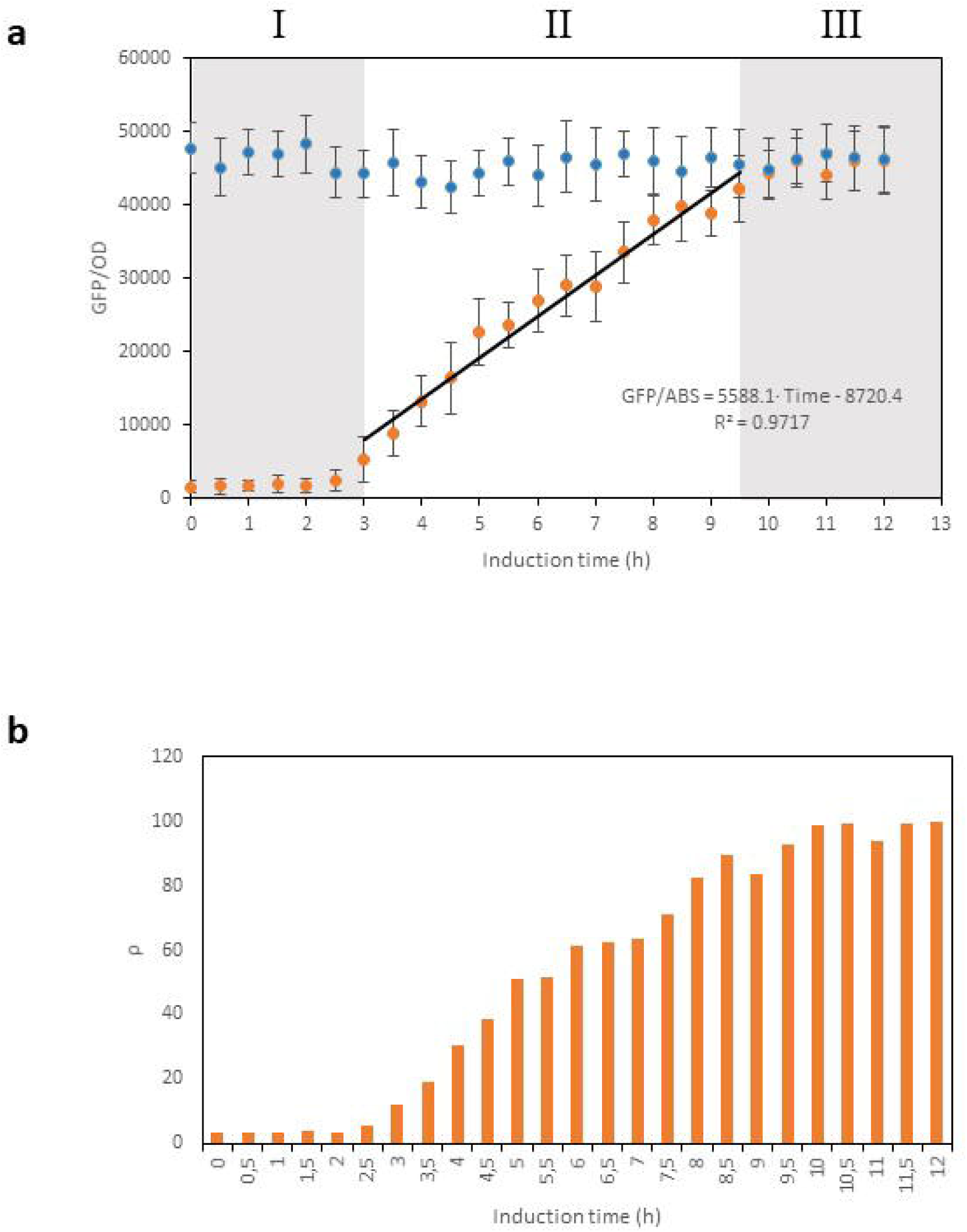
a. Relationship between the induction time of Bxb1 expression and resultant GFP expression levels following recombination mediated by Bxb1 using cell types C1 (orange points) and C2 (blue points). While GFP expression levels are independent of the Bxb1 levels in cell type C2 because GFP expression is not dependent on Bxb1, the GFP expression levels are dependent on the level of Bxb1 induction in cell type C1, with three distinct behavioral regions. In Region I, no significant GFP expression is observed, indicating that intracellular Bxb1 levels are insufficient to trigger gene expression as a result of recombination events. In Region II, a quasi-linear relationship is observed between the Bxb1 induction time and GFP expression level (r²=0.97). GFP expression levels peak in Region III, in which they are equal to those in cell type C2. **b.** Percentage of GFP expression in cell type C1, i.e., ρ, relative to the maximum levels in cell type C2. Experimental values represent the mean and error bars indicate the standard deviation from three independent experiments.

These results can be compared with GFP expression levels resulting from recombination events in cultures of cell type C1 that were exposed to arabinose throughout exponential phase. Figure S3 displays GFP expression levels upon addition of different concentrations of arabinose. GFP expression levels were significantly lower than those observed upon temporal induction of Bxb1 in the same cell type. While continuous expression of Bxb1 can offset the dilution effects associated with cellular duplication, it seems the continuous Bxb1 expression likely has adverse effects on GFP expression levels compared with temporal Bxb1 induction.

These results indicate that gene expression can be precisely controlled by applying temporal pulses of external signals in a given temporal range.

### Effect of the duration of exponential phase on gene expression mediated by Bxb1

The previous results demonstrated that the recombination efficiency and level of intracellularly accumulated Bxb1 accurately determine the recombination level and consequently associated gene expression. However, the duration of exponential phase is crucial for gene expression and consequently can affect the recombination efficiency (Figure S1). To evaluate this effect, Bxb1 expression was induced for 5 h in a culture of cell type C1, after which the culture entered stationary phase and plasmid gene expression significantly declined. Specifically, a saturated culture of cell type C1 lacking arabinose was prepared and divided into two equal volumes. The first sample was centrifuged and the supernatant was collected and passed through a 0.22 µm filter to ensure all cells were removed. This procedure resulted in a culture medium lacking sufficient nutrients and free of cells and arabinose (see Material and Methods for details).

The second sample was diluted by adding fresh LB to an OD of 0.45, and 10^-4^ M arabinose was added to induce Bxb1 expression. After 5 h of growth at 37°C, the culture had re-entered stationary phase. At this point, the pellet was separated from the supernatant by centrifugation and resuspended in the arabinose-free supernatant obtained from the first sample. This produced a cell type C1 culture that had been exposed to arabinose for 5 h during exponential phase, meaning Bxb1 was induced during this time period, and was subsequently maintained in stationary state in medium without nutrients available and lacking arabinose.

Finally, a series of cultures was prepared by diluting the stationary phase culture with intracellular Bxb1 into fresh medium at various concentrations. This yielded a set of cultures with the same intracellular Bxb1 level but different initial concentrations. These cultures were cultured overnight at 37°C with shaking. Figure 3 shows the GFP expression levels resulting from recombination (orange bars) and the corresponding recombination efficiency ρ (blue line). Although all cultures initially had the same intracellular concentration of Bxb1 and no more Bxb1 was produced during the growth period due to the absence of arabinose, GFP expression levels differed depending on the initial concentration of the culture. Considering the relationship between the initial culture concentration and duration of exponential phase (Figure S2), these differences indicate that cultures with higher initial concentrations have a shorter exponential phase, which results in lower GFP expression due to the culture entering stationary phase. As the initial concentration of the culture decreased and thus the duration of exponential phase increased, GFP expression levels rose, reaching a ρ value of 100%.

**Figure 3.**
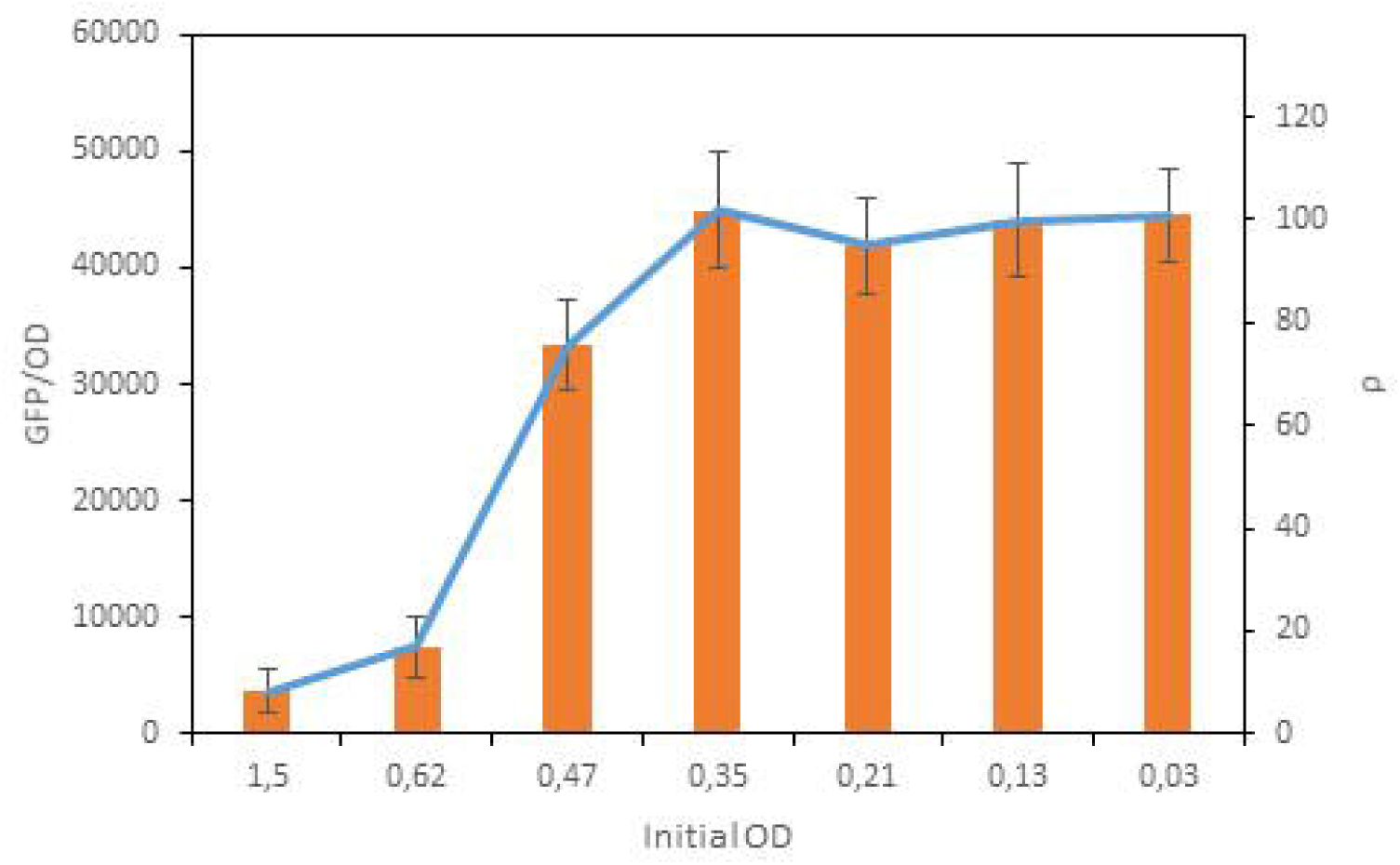
Gene expression levels mediated by recombination. The orange bars display GFP expression levels measured in various cultures of cell type C1. Although the intracellular Bxb1 level is the same in all cultures, depending on the initial concentration of the culture, the GFP expression levels vary considerably due to the differing durations of exponential phase. The blue line represents the efficiency of gene expression (ρ) mediated by recombination events. Experimental values represent the mean and error bars indicate the standard deviation from three independent experiments.

However, these results do not clarify the cause of the variations in GFP expression levels. This variation could be due to differences in recombination efficiency, specifically the number of sites recombined, which may be influenced as the culture transitions into stationary phase. Alternatively, recombination events might occur, but expression of the reporter protein, i.e. GFP, may not be feasible in stationary phase.

### Analysis of Bxb1 activity during stationary phase

To determine whether Bxb1 activity is maintained during stationary phase, although recombination during this phase does not increase GFP expression, an assay was conducted under the previously described conditions. A saturated culture of cell type C1 without arabinose was diluted to achieve an OD of 0.45 and then 10^-4^ M arabinose was added. The culture was maintained at 37°C for 5 h until it re-entered stationary phase. At this point, the pellet was separated from the supernatant containing arabinose and resuspended in the original, arabinose-free supernatant from the saturated culture to maintain cells in stationary phase.

Cells in this new stationary phase contained a pool of intracellular Bxb1 produced during the 5 h of induction with arabinose. The culture was then incubated at 37°C for 20 h. Periodically, 1 µL of this culture was harvested and diluted in 5 mL of fresh LB medium lacking arabinose. These newly diluted cultures were maintained at 37°C overnight before GFP expression levels were measured.

If Bxb1 retained its activity during stationary phase, the number of recombined sites would increase over time, despite GFP expression levels remaining stable. However, upon addition of fresh medium, cells would re-enter exponential phase, in which GFP expression would depend on the number of recombined sites. Conversely, if Bxb1 activity was limited by entry into stationary phase, the GFP expression levels would remain stable across all cell cultures.

Figure 4a displays the GFP expression levels in different cultures. The blue bars represent GFP expression levels in the stationary culture measured at various time points after a 1 µL inoculum was taken. The orange bars represent GFP expression levels in the resulting culture, which was obtained by diluting this inoculum in fresh, arabinose-free medium and then incubated overnight at 37°C. While GFP expression levels, which stabilized at ρ=18%, only slightly increased in the stationary culture over time, they increased significantly, up to ρ=100%, when the culture re-entered exponential phase as a result of dilution in fresh medium. These time-dependent changes of GFP expression levels corresponded with the number of recombined sites, which increased over time during the stationary period. These results indicate that Bxb1 activity is maintained during stationary phase, despite the inability to monitor this activity through GFP expression. After 20 h in stationary phase, the recombination efficiency reached ρ=100%. While induction of Bxb1 for 5 h yielded a recombination efficiency of ρ=51% (Figure 2b), the same induction followed by entry into stationary phase yielded a recombination efficiency of ρ=100%. This is explained by the fact that recombination events were occurring during cellular growth in Figure 2b, which implies that the concentration of intracellular Bxb1 decreases upon cellular division because cells do not produce more Bxb1 due to the absence of arabinose. However, when the culture enters stationary phase after Bxb1 induction, this dilution effect associated with cellular division does not occur, meaning the concentration of intracellular Bxb1 only decreases due to proteolytic degradation, which takes much longer. Consequently, maintenance of the intracellular pool of Bxb1 ensures a much higher recombination efficiency, which reaches 100%. These results suggest that maintaining recombinase activity during stationary phase is more efficient than maintaining recombinase activity during exponential phase.

**Figure 4.**
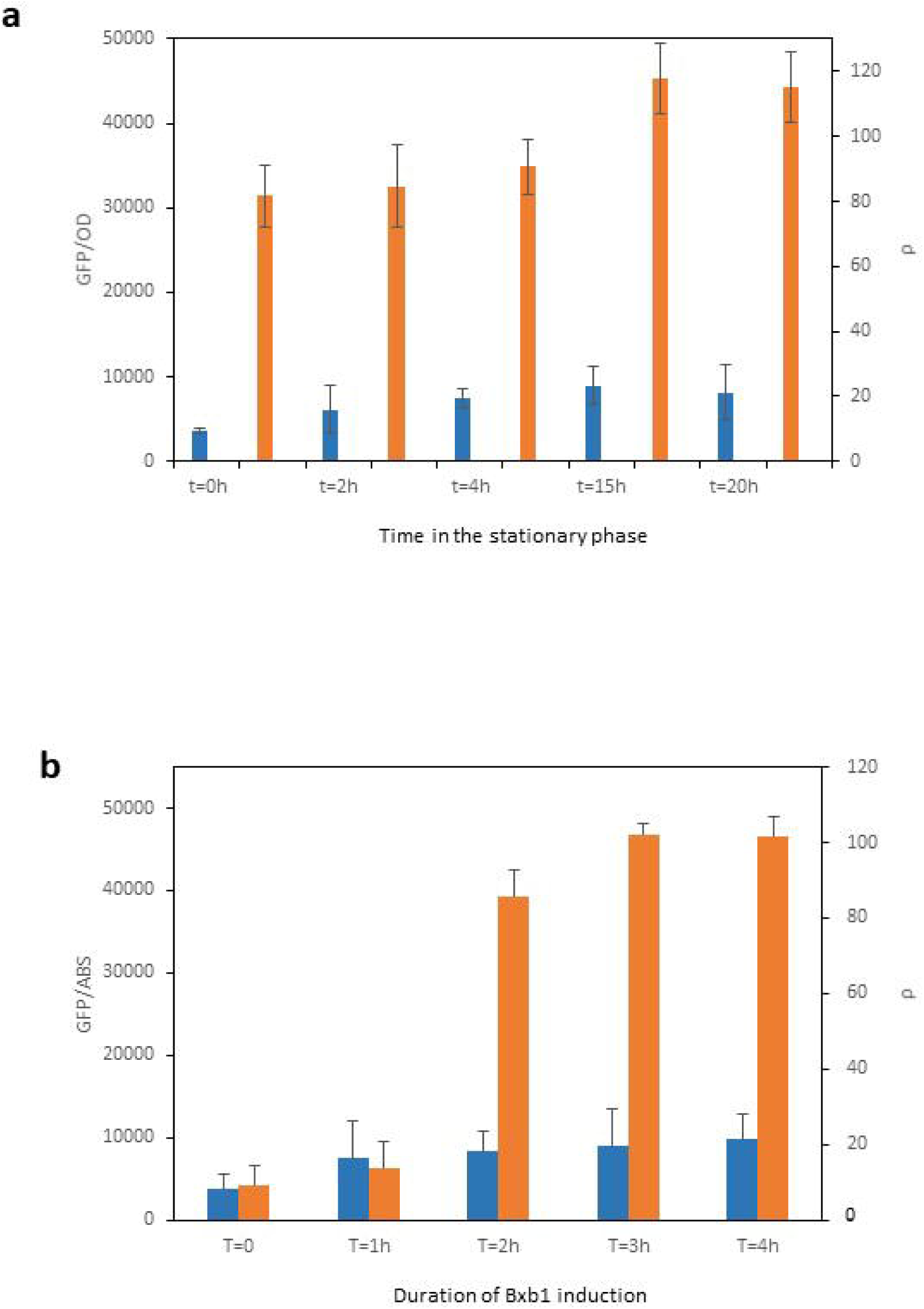
Gene expression mediated by recombination during stationary phase. **a.** Blue bars represent the GFP expression levels measured in a culture of cell type C1 after Bxb1 induction for 5 h with different durations in stationary phase. Orange bars represent the same culture after identical durations in stationary phase, after which the culture was returned to exponential phase via dilution in fresh medium. The difference in GFP expression upon return to exponential phase, compared with that in the corresponding culture in stationary phase, indicates that Bxb1 activity is maintained during stationary phase, although no increase in GFP expression levels is observed. However, when the culture re-enters exponential phase, GFP is expressed, with higher GFP expression levels correlating with longer durations of stationary phase. The right axis indicates the gene expression efficiency, ρ. **b.** Correlation between gene expression levels and time of Bxb1 induction in cultures maintained in stationary phase (blue bars) and in the same cultures after re-entering exponential phase (orange bars). The right axis indicates the gene expression efficiency, ρ. Experimental values represent the mean and error bars indicate the standard deviation from three independent experiments.

Subsequently, the minimum induction period of Bxb1 prior to entry into stationary phase that achieves the maximal recombination efficiency was explored. The same methodology was applied but different dilutions of the initial saturated culture were used, corresponding to various periods before entry into stationary phase. The lower the dilution, the shorter the Bxb1 induction time. Cultures were adjusted to have exponential phases of 1, 2, 3, and 4 h. After these periods, arabinose was removed from the cultures, which were then suspended in the arabinose-free supernatant from the original saturated culture to ensure that the culture remained in stationary phase. Figure 4b displays the results. Blue bars represent GFP expression levels in stationary phase of each culture, whereas orange bars represent GFP expression levels in cultures prepared by taking a 1 µL inoculum from the corresponding stationary cultures and diluting them in 5 mL of arabinose-free LB medium, which were then incubated overnight at 37°C.

While the culture maintained in stationary phase exhibited slight increases in GFP expression levels and ρ values (blue bars in Figure 4b), markedly different behavior was noted when cultures were diluted, i.e., they re-entered exponential phase. Specifically, although 1 h of Bxb1 induction was insufficient to induce significant recombination events, 2 and 3 h of Bxb1 induction achieved a recombination efficiency of 86% and 100%, respectively. The same amounts of induction followed by exponential phase resulted in much lower recombination efficiencies (Figure 2a, b). These results indicate that while Bxb1 activity does not seem to be limited by entry into stationary phase, gene expression of plasmids is affected. This scenario allows more efficient recombination because the Bxb1 concentration is not reduced by cell division during stationary phase. Consequently, higher recombination efficiencies can be achieved with a stable Bxb1 concentration.

Finally, recovery of gene expression of recombined plasmids by adding nutrients to the saturated medium instead of diluting the culture in fresh medium was explored. This approach would allow the recovery of gene expression from plasmids without reducing the cellular concentration, which could be beneficial in many applications. In this case, similar cultures of C1 were prepared. Each culture was prepared from a saturated culture of cell type C1 without arabinose, diluted to achieve an OD of 0.45 with the addition of 10⁻⁴ M arabinose. Each culture was maintained at 37°C for 5 hours until they re-entered the stationary phase. At this point, the pellet was separated from the supernatant containing arabinose and resuspended in the original arabinose-free supernatant from the saturated culture to maintain cells in the stationary phase. At 20 h after the removal of arabinose, powdered nutrients were added to these stationary phase cultures according to the compositions shown in Table 1. After the addition of nutrients, cultures were maintained at 37°C overnight. The addition of nutrients greatly increased GFP expression levels (Figure 5).GFP expression levels varied according to the concentration of nutrients added; the more nutrients added, the higher the recombination efficiency. These results reinforce the idea that Bxb1 remains active during stationary phase and that gene expression is primarily perturbed by depletion of key nutrients.

**Figure 5.**
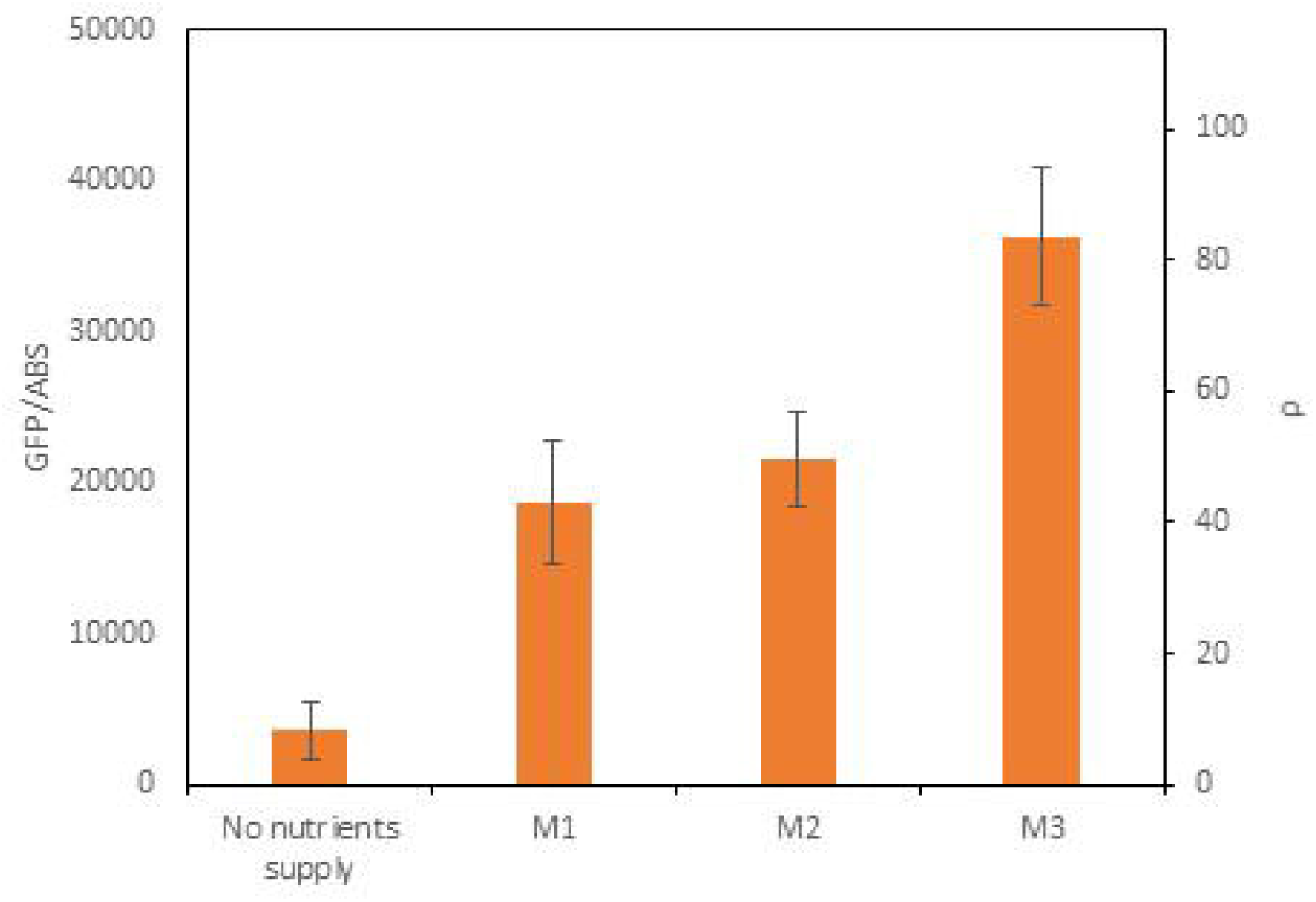
GFP expression levels in a culture of cell type C1 after Bxb1 induction for 5 h followed by entry into stationary phase according to the amount of nutrients supplied. Following nutrient addition, GFP expression resulting from recombination events occurs. Depending on the quantity of nutrients available, cells have more or less time to express GFP, resulting in different levels of fluorescence. The right axis indicates the gene expression efficiency, ρ. Experimental values represent the mean and error bars indicate the standard deviation from three independent experiments.

**Table 1.** Nutrients supplied to cell cultures.

|  |  |
| --- | --- |
| M1 | 5 g/L tryptone<br>2.5 g/L yeast extract |
| M2 | 10 g/L tryptone<br>5 g/L yeast extract |
| M3 | 20 g/L tryptone<br>10 g/L yeast extract |

## MATERIALS AND METHODS

### Strains, media, and growth conditions

*E. coli* TOP10 (Invitrogen, USA) was used in all cloning and expression experiments. Cells were grown in LB medium at 37°C with shaking at 200 rpm and selected with appropriate antibiotics (35 µg/mL chloramphenicol or kanamycin; Sigma, USA). Bacterial strains were preserved in LB medium containing 20% (v/v) glycerol at −80°C.

All culture were preprared from single colonies obtained from streaked glycerol stocks were inoculated into fresh LB medium containing antibiotics and grown overnight at 37°C with shaking at 200 rpm. Induction medium was LB medium supplemented with the appropriate antibiotic and L-arabinose (Sigma).

All results are the means of three independent experiments. Error bars represent the standard deviation of these experiments.

### Building genetic circuits

All genetic circuits were constructed using the Biobrick assembly method (Ginkgo Bioworks, USA). The constructs were built using one of the following backbones: pSB1AC3 (high-copy plasmid with ampicillin and chloramphenicol resistance) and pSB3K3 (low-copy number plasmid with kanamycin resistance). All transformations were performed using a chemical method, and the sequences of genetic constructs were confirmed by Sanger sequencing.

### Fluorescence quantification in liquid cultures

GFP fluorescence was measured at an excitation wavelength of 485 ± 20 nm and an emission wavelength of 528 ± 20 nm using a Synergy MX microplate reader (BioTek Instruments, USA) and the absorbance was measured at wavelength of 600nm. Sample S absorbance termed OD600 (S), and fluorescence, termed f (S), were corrected using the respective background signals of controls, namely, OD600 (B) and f (B). Reporter protein Θ fluorescence was calculated using the following equation:

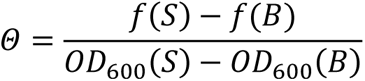

### Experiments characterizing the effects of the culture growth phase on plasmid gene expression

Single colonies of cell type C3 obtained from streaked glycerol stocks were inoculated into 10 mL of fresh LB medium containing kanamycin and grown overnight at 37°C with shaking at 200 rpm. The next day, 13 cultures were prepared with different concentrations of the initial inoculum obtained from the saturated culture. This was achieved by performing 9:10 serial dilutions (mixing 3960 µL of the previous dilution with 40 µL of fresh LB medium). All cultures were supplemented with 10^-4^ M arabinose. These cultures were grown overnight at 37°C with shaking at 200 rpm.

### Experiments characterizing the effect of the Bxb1 expression level on the recombination efficiency

Single colonies of cell types C1 and C2 obtained from streaked glycerol stocks were inoculated into 5 mL of fresh LB medium containing kanamycin and chloramphenicol and grown overnight at 37°C with shaking at 200 rpm.

The next day, from the saturated cultures of cell types C1 and C2, new cultures were prepared by adding 40 µL of each culture to 3960 µL of fresh LB medium containing kanamycin and chloramphenicol. These cultures were supplemented with 10-4 M arabinose and maintained at 37°C with shaking at 200 rpm.

Every 30 min, a 1 mL sample of each culture was taken and centrifuged to separate the pellet from the supernatant. The pellet was resuspended in the same volume of fresh LB medium lacking arabinose. Finally, 1 µL of this resuspended pellet was used to inoculate 5 mL of LB medium lacking arabinose. These cultures were maintained overnight at 37°C with shaking at 200 rpm.

### Experiments characterizing the effect of the duration of exponential phase on gene expression mediated by Bxb1

Single colonies of cell type C1 obtained from streaked glycerol stocks were inoculated into 40 mL of LB medium containing kanamycin and chloramphenicol and grown overnight at 37°C with shaking at 200 rpm. This culture was divided into two equal volumes of 20 mL. Each volume was centrifuged for 10 min at 14,000 rpm, and the supernatant was collected and filtered through a 0.22 µm Millipore® filter to ensure all cells were removed.

One volume of the culture was diluted by adding fresh LB medium to achieve an OD of 0.45, supplemented with 10-4 M arabinose to induce Bxb1 expression, and maintained at 37°C with shaking at 200 rpm. Periodically, OD600 was measured until the culture re-entered stationary phase. At this point, the pellet was separated from the supernatant by centrifugation at 14,000 rpm for 10 min and resuspended in the arabinose-free supernatant obtained from the other volume of the culture.

Finally, a series of new cultures was prepared at different dilutions. All these cultures were grown overnight at 37°C with shaking at 200 rpm.

### Addition of powdered nutrients to cultures in stationary phase

Single colonies of cell type C1 obtained from streaked glycerol stocks were inoculated into 40 mL of LB medium containing kanamycin and chloramphenicol and grown overnight at 37°C with shaking at 200 rpm. Once the culture had been in stationary phase for 20 h, powdered nutrients, specifically tryptone and yeast extract, were added. Several experiments were conducted by adding different amounts of these nutrients as described in Table 1. After addition of nutrients, the cultures were maintained overnight at 37°C with shaking at 200 rpm.

## CONCLUSIONS

The use of recombinases capable of performing DNA modifications in response to external stimuli holds great potential in the field of synthetic biology for the development of complex genetic circuits with multiple applications. This is because such recombinases can be reconfigured in vivo in response to specific inputs, thus altering cellular behavior in a controlled manner. In many of these applications, the recombination efficiency, i.e., the proportion of recombined sites relative to all possible recombination sites, must be precisely controlled. For instance, in some applications, partial recombination, i.e., not all possible sites are recombined, can lead to incorrect behaviors in genetic circuits due to the coexistence of two populations of plasmids, i.e., those that have been recombined and those that have not. In other cases, however, it may be desirable to induce partial recombination in a controlled manner. Understanding the parameters that affect the recombination efficiency is crucial to design biological systems that respond predictably and controllably.

A notable aspect is that the recombination efficiency depends not only on the induction level of the corresponding recombinase but also on external factors, such as the growth phase of the cellular culture. This study shows that the recombination efficiency can be precisely controlled by controlling the duration that cells are exposed to the external input responsible for inducing expression of the recombinase. However, the phase of the cell culture when the external input is introduced markedly affects gene expression regulated by recombination events. When cultures are exposed to the external inducer during exponential phase, gene expression regulated by recombination correlates with the recombination efficiency. Specifically, in the case of Bxb1, there is a range of recombinase induction times that correlate quasi-linearly with resultant gene expression. This means it is possible to precisely establish how long the genetic circuit should be exposed to the external inducer, e.g., arabinose, in order to achieve recombination in a proportion of the entire population of plasmids susceptible to recombination. Temporal induction of Bxb1 expression during exponential phase yields better results in terms of GFP expression than sustained induction of Bxb1 expression throughout this phase.

However, if the time required for the cell culture to reach stationary phase is shorter than the time required to perform recombination, resultant gene expression stops correlating with the recombination efficiency because gene expression from plasmids decreases during stationary phase. Consequently, not only the level of recombinase expression, but also the timing of when the culture is exposed to the external input is a key parameter to ensure that gene expression correctly correlates with the recombination efficiency.

When the culture enters stationary state, although observed gene expression does not correspond with the population of recombined plasmids, recombinase activity is maintained. Consequently, if after a period in stationary state the culture re-enters a phase that allows gene expression of plasmids, all recombination events performed surreptitiously during stationary phase will increase gene expression. Given there is no significant cellular division during stationary phase, this allows the recombination efficiency to reach 100% with lower levels of the intracellular recombinase than are necessary if the recombination events occur during exponential phase.

This suggests a strategy to maximize the effect of recombination by inducing expression of recombinases shortly before entry into stationary phase. Specifically, in the case of Bxb1, exposing the culture to the external input 2 and 3 h before entry into stationary phase achieved a recombination efficiency of 86% and 100%, respectively. When the same levels of Bxb1 expression were induced during stationary phase, the recombination efficiencies were 3.4% and 12%.

However, translating the recombination events performed during stationary phase into corresponding gene expression requires modification of the cell culture conditions. One option is to dilute the culture in fresh medium. In this case, the levels of gene expression achieved depend on the degree of dilution. Therefore, to ensure that the maximum levels of gene expression are achieved, it is advisable to perform large dilutions in order to maximize the duration that the culture remains in exponential phase.

In certain applications, both industrial and biomedical, dilution of cultures is undesirable or impossible. In this case, gene expression of plasmids can be recovered by adding nutrients directly to the culture in stationary state. This provides a simple way to control gene expression regulated by recombination events and indicates that the reduction in gene expression of plasmids during stationary phase is directly associated with depletion of key nutrients present in the culture medium, meaning that supplementing the medium with these nutrients allows the gene expression machinery to be recovered.

Although the results help to understand and design strategies for genetic regulation by recombinases, this study has several limitations. First, *E. coli* was used as a model organism and Bxb1 was used as a model recombinase. Although *E. coli* and Bxb1 are very common elements in the context of synthetic biology, a broader study is necessary using other organisms and recombinases to analyze the generality of the results. Second, this study focused on one type of recombination, i.e., DNA excisions. It is necessary to extend this analysis to other types of recombination in order to establish the generality of the results. Third, the genetic constructs used in this study were integrated into plasmids. Plasmids are very commonly used for the development and validation of experimental genetic circuits due to their simplicity and efficacy. However, in applications that require the stability of genetic constructs to be ensured, such constructs are usually integrated into the cellular genome. Consequently, it is necessary to perform experiments in integrated genetic systems in order to analyze the effects of the level of recombinase induction and the growth phase of the cell culture on the recombination efficiency and associated gene expression.

## DATA AVAILABILITY STATEMENT

The raw data supporting the conclusions of this article will be made available by the authors, without undue reservation.

## ACKNOWLEDGEMENTS

We thank all the members of the Synthetic Biology for Biomedical Applications Laboratory for fruitful conversations.

## AUTHOR CONTRIBUTIONS

M.G.-C. and J.M. designed the circuits. M.G.-C. and J.M. performed the experiments. J.M. and M.G.-C. wrote the paper.

## SUPPLEMENTARY MATERIAL

The Supplementary Material for this article can be found online at:

