## Supplemental Figures for "CHARACTERIZATION OF RECOMBINASE ACTIVITY ACROSS CELLULAR GROWTH PHASES"

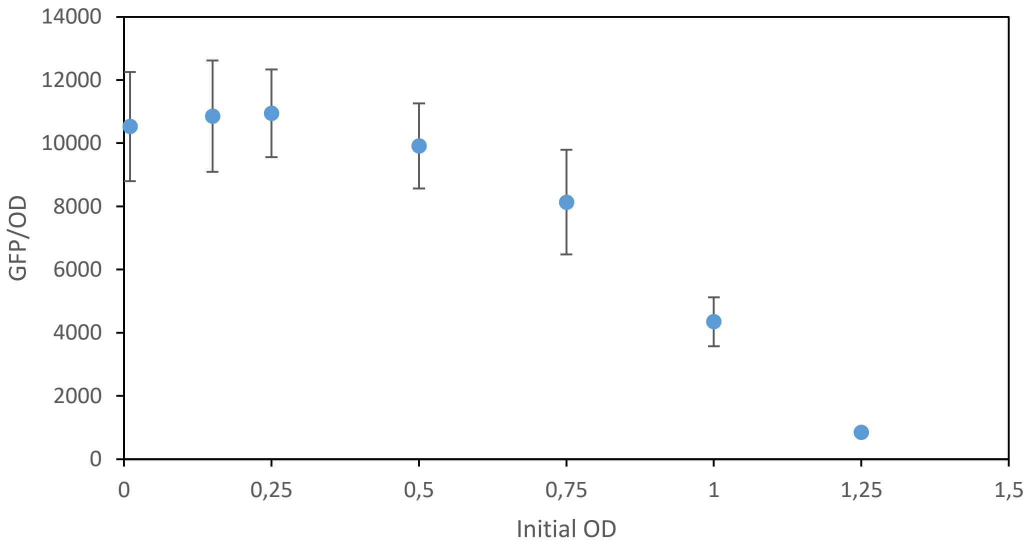


**Figure S1 .** GFP expression levels in cultures of cell type C3 induced by $10^4$ M arabinose according to the initial concentration of the culture. Although the concentration of arabinose is the same in all cultures, GFP expression levels depend on the initial concentration of the culture. Higher initial culture concentrations result in lower GFP expression levels. Experimental values represent the mean and error bars indicate the standard deviation from three independent experiments.

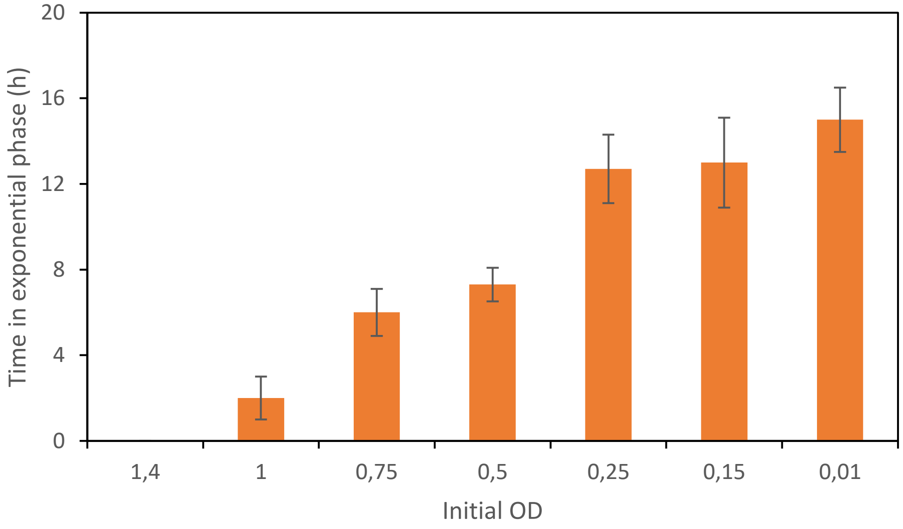


**Figure S2.** Relationship between the duration of exponential phase in a culture of cell type C3 and the initial concentration of the culture. Higher initial concentrations of the culture result in shorter exponential phases. Experimental values represent the mean and error bars indicate the standard deviation from three independent experiments.

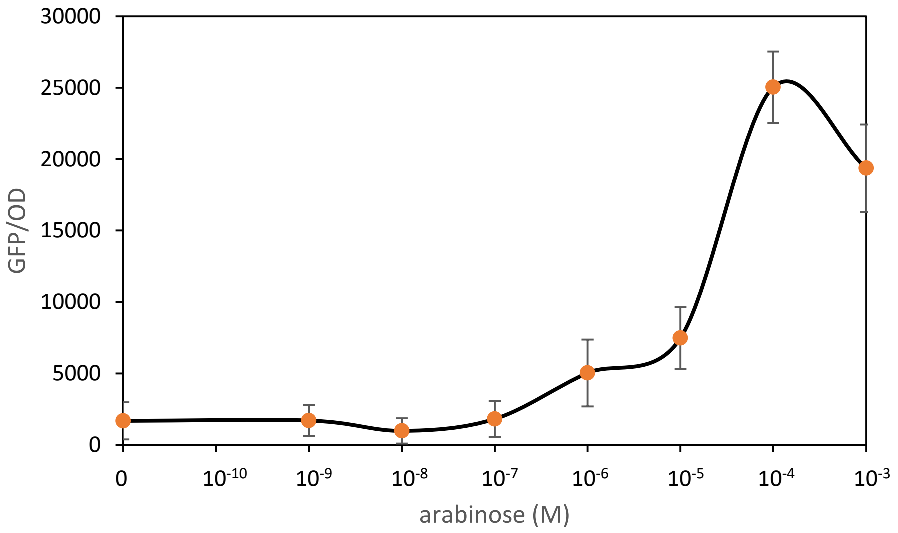


**Figure S3.** Relationship between the arabinose concentration and GFP expression mediated by recombination events. Experimental values represent the mean and error bars indicate the standard deviation from three independent experiments.
